# Sparkling Water Co-Consumption Enhances Oxytocin-Linked HR Synchrony and Social Connection in Live Sports Spectatorship

**DOI:** 10.64898/2026.01.05.697802

**Authors:** Takashi Matsui, Takafumi Yamaguchi, Shohei Dobashi, Shion Takahashi, Yukina Tachibana, Ren Takamizawa, Wataru Kosugi, Seiichi Mizuno

## Abstract

Live sport spectatorship can promote social connection, yet scalable, low-burden strategies to strengthen bonding in real-world spectator settings remain limited. We here tested the hypothesis that co-consuming sparkling water during spectating attenuates hunger and fatigue and strengthens enjoyment and perceived unity, potentially via HR synchrony linked to oxytocin dynamics. In an individually randomized field experiment at a collegiate women’s basketball home game (sparkling water: n = 21; plain water: n = 19), spectators consumed 200 mL of sparkling water or plain water in synchrony before tip-off and before the end of halftime, with additional ad libitum intake of the assigned beverage permitted thereafter. We assessed acute freshness and hunger, repeated fatigue and enjoyment, interpersonal HR synchrony from the first half to 15-min post-game, perceived unity toward players and fellow spectators, and salivary oxytocin levels. Relative to plain water, sparkling water increased freshness and reduced hunger after intake, attenuated post-game fatigue, and enhanced enjoyment and perceived unity. HR synchrony was preserved in association with attenuated post-game fatigue after the game, and sustained synchrony was associated with higher perceived unity. Oxytocin decreased more in the sparkling-water group, and this decrease was associated with preserved HR synchrony as well as with changes in unity toward fellow fans. These findings support our hypothesis and suggest that co-consuming sparkling water serves as an alcohol-free, low-burden ritual to enhance social connection during and after live sport events.

**Highlights:**

- Sparkling water increased freshness and reduced hunger after intake in spectators.
- Sparkling water attenuated the post-game increase in subjective fatigue.
- Sparkling water increased enjoyment and perceived unity.
- Post-game HR synchrony was preserved in association with lower fatigue and higher unity.
- Oxytocin decreased more with sparkling water and tracked HR synchrony and fan unity.
- Co-consuming sparkling water is a low-burden, alcohol-free ritual for social connection.

## 1. Introduction

Live sport spectatorship is a naturalistic collective emotional experience that can strengthen social connection by aligning attention, affect, and identity among individuals who share the same event in real time (Inoue et al., 2022). Such group experiences are increasingly recognized as behaviorally meaningful, with evidence linking attendance at live sporting events to higher subjective well-being and lower loneliness (Keyes et al., 2023). Consistent with this view, longitudinal studies in Japan suggest that sports watching is associated with improved mental and social well-being, including fewer depressive symptoms and richer social connections (Kawakami et al., 2024; Tsuji et al., 2026).

A central challenge is to specify the socio-biological processes through which spectatorship becomes felt social connection. One candidate pathway is interpersonal physiological synchrony, defined as temporal alignment (including lagged covariation) in autonomic signals such as heart rate (HR) across individuals (Palumbo et al., 2017). Oxytocin represents another candidate mechanism given its established role in affiliative motivation and bonding. In our recent field evidence, live sport spectating engaged endogenous oxytocin-related dynamics together with HR synchrony, consistent with a socio-physiological route to social connection (Matsui et al., 2025). In the same line of work, stronger HR synchrony was linked to greater perceived unity toward both players and fellow spectators, assessed using the Inclusion of Other in the Self (IOS) scale. These findings position the synchrony-to-IOS pathway as a measurable bonding-relevant route in a real-world collective setting (Matsui et al., 2025), and motivate a practical next question: can this pathway be causally strengthened in the field by a scalable, low-burden intervention?

Despite progress in identifying candidate mechanisms, causal and scalable strategies to strengthen bonding-related outcomes in spectator settings remain limited, particularly approaches that target coupling among spectators and downstream unity. A plausible intervention is a simple co-consumption ritual. In humans, incidental similarity in food consumption increases interpersonal trust and cooperation (Woolley & Fishbach, 2017), and in wild chimpanzees, food sharing has been linked to higher oxytocin responses than grooming (Wittig et al., 2014). Together, these findings suggest that sharing the same drink could serve as a repeatable add-on capable of amplifying socio-physiological coupling and perceived unity during live sport spectatorship.

Matchday consumption rituals in real venues are often alcohol-centered. Alcohol availability in sporting venues has been discussed in relation to alcohol-related problems and venue safety or public health concerns (Ruehlmann et al., 2023; Lenk et al., 2010; Nelson & Wechsler, 2003). Accordingly, growing interest has emerged in appealing alcohol-free options that preserve, and potentially amplify, the social benefits of collective events without relying on alcohol-centered rituals (Buss et al., 2025; Waehning & Wells, 2024; Yoshimoto et al., 2023).

From this perspective, an ideal beverage-based intervention should be alcohol-free, appealing enough to function as a shared matchday ritual, and capable of modulating subjective state to support sustained engagement during and after the game. Sparkling water (carbonated water) is a suitable candidate because widespread availability as a non-alcoholic option combines with distinctive oral sensory stimulation that can influence affective and neurophysiological states relevant to engagement. In our recent work on prolonged esports play, sparkling water reduced subjective fatigue and hunger and increased enjoyment (Takahashi et al., 2025). In addition, carbonated water ingestion increased oxyhemoglobin in a left frontal site measured with near-infrared spectroscopy, located near the orbitofrontal cortex and implicated in reward valuation processes (Kosugi et al., 2024).

We therefore hypothesized that co-consuming sparkling water attenuates hunger and fatigue and enhances enjoyment and perceived unity, potentially via post-game HR synchrony linked to salivary oxytocin dynamics. To test this hypothesis, we conducted an individually randomized field experiment comparing the sparkling-water group (SW) with the plain-water group (PW) during a collegiate women’s basketball home game, assessing subjective state, IOS-based perceived unity, HR synchrony, and salivary oxytocin.

## 2. Materials and Methods

### 2.1 Study design and setting

We conducted a naturalistic, individually randomized, parallel-group field experiment to test whether co-consuming sparkling water during live sport spectatorship strengthens socio-physiological coupling and perceived unity. The study was implemented at a Japanese collegiate women’s basketball home game held at the Central Gymnasium, University of Tsukuba (Japan) in August 2025. All participants attended the same game and completed the study procedures during a single spectating session. The experimental design, standardized beverage intake, and assessment timeline are summarized in Figure 1. Ethical approval for this study was granted by the Research Ethics Board of the University of Tsukuba, and all procedures complied with the Declaration of Helsinki.

**Figure 1.**
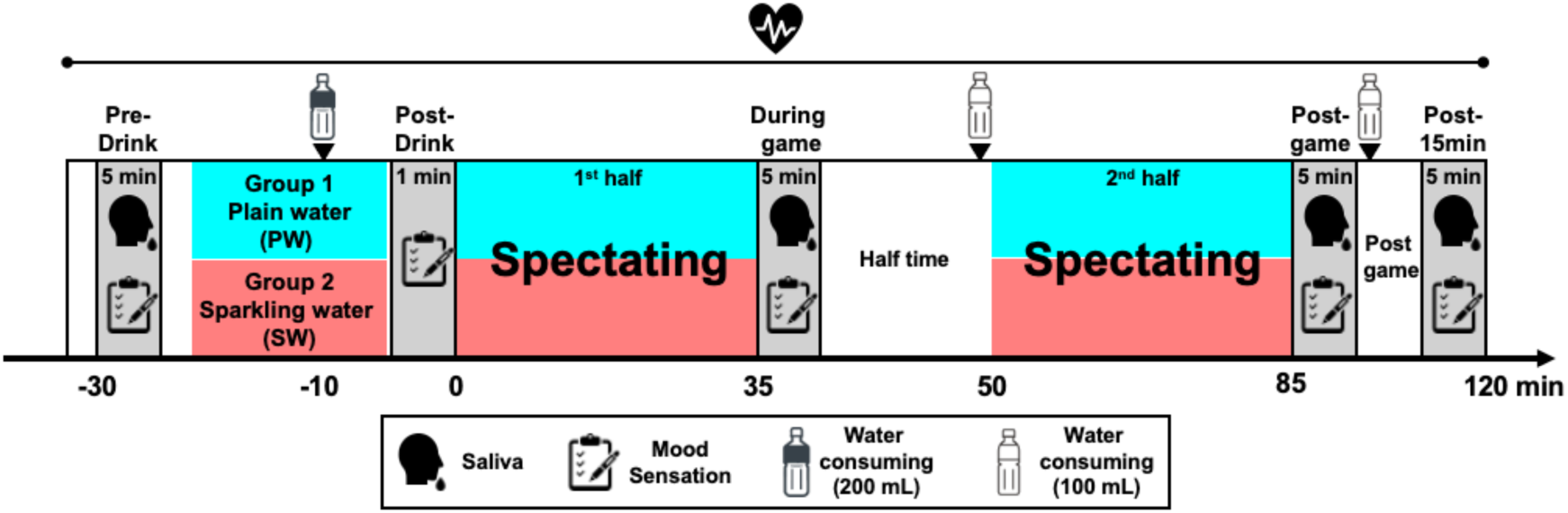
Experimental design and assessment timeline. Schematic of the individually randomized field experiment during a collegiate women’s basketball home game. Spectators were assigned to plain water (PW) or sparkling water (SW). The timeline indicates standardized beverage consumption (200 mL immediately before tip-off and immediately before the end of halftime), acute pre–post assessments around the first intake, repeated assessments (pre-game, halftime/during game, immediately post-game, and 15 min post-game), and saliva collection, with heart rate recorded continuously.

### 2.2 Participants and recruitment

Participants were spectators recruited via an online ticketing website. Individuals purchased a “research-participation ticket” priced at JPY 2,000, which included a JPY 1,500 honorarium upon completion of the study procedures. Exclusion criteria were a history of cardiovascular disease, current medication use, pregnancy, smoking, and insufficient Japanese proficiency. Participants were instructed to refrain from alcohol consumption and strenuous exercise from the day before through the day of the experiment, to avoid caffeine intake on the day of the experiment, and to complete their last meal at least 2 h before game start.

An a priori power analysis was conducted using G*Power (version 3.1) for a two-tailed independent-samples t-test to approximate the primary between-group comparison. Assuming a large effect size (Cohen’s d = 0.9), α = 0.05, and power (1 − β) = 0.80, the required sample size was N = 42 (21 per group). We therefore aimed to recruit approximately 40 participants.

The final analyzed sample comprised 40 spectators (PW: n = 19; SW: n = 21). All participants provided written informed consent prior to participation. Participant characteristics are summarized in Table 1.

**Table. 1.**
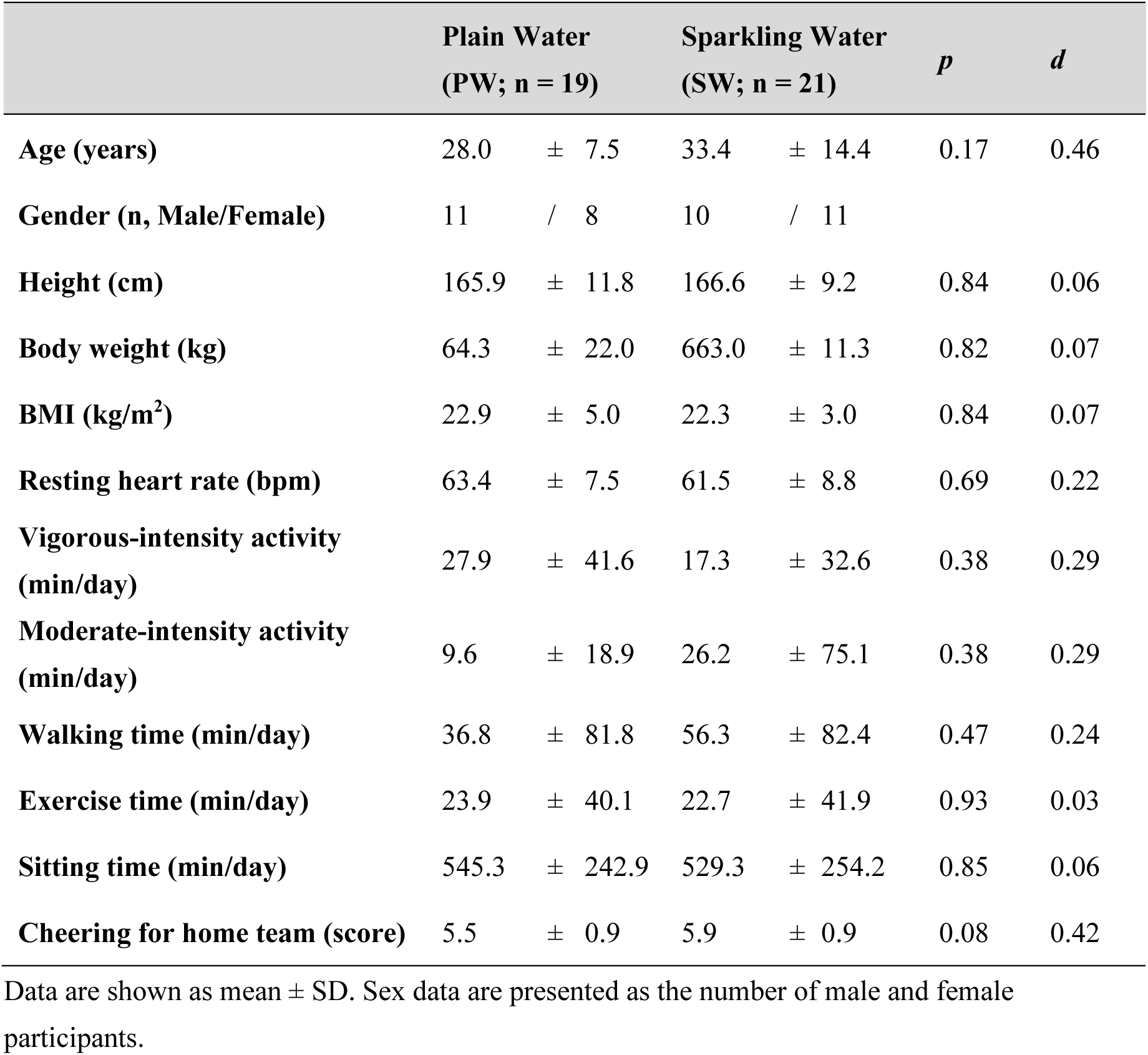
Participants’ characteristics.

### 2.3 Randomization, allocation, and blinding

Participants were individually randomized to SW or PW groups using a random number table with stratification by sex and age decade (teens, 20s, 30s, 40s, 50s, and 60s) to balance these characteristics across groups. The stratified randomization sequence was prepared in advance and implemented on-site on the day of the game; participants were assigned to a group at enrollment according to the pre-generated allocation list.

Because the intervention involved beverages with discernible sensory properties, participant blinding was not feasible. All participants received standardized instructions and underwent identical procedures across groups, except for the assigned beverage.

### 2.4 Intervention

Participants in the SW group were provided with commercially available sparkling water (Wilkinson; Asahi Soft Drinks Co., Ltd., Japan), whereas participants in the PW group were provided with plain water. Beverages were stored refrigerated and taken out immediately before distribution.

To standardize co-consumption timing, participants consumed 200 mL of their assigned beverage synchronously at two predefined time points: immediately before tip-off and immediately before the end of halftime. After these standardized intakes, participants were allowed to drink the assigned beverage ad libitum for the remainder of the session. Participants were instructed to consume only the assigned beverage during the experiment. Ad libitum intake volume was not quantitatively recorded.

### 2.5 Procedures and timeline

Upon arrival at the venue, participants provided informed consent and received standardized instructions regarding pre-experiment restrictions (alcohol and strenuous exercise from the day before through the day of the experiment; no caffeine on the day of the experiment; last meal completed ≥2 h before game start). Participants were then enrolled and assigned to SW or PW groups according to the stratified randomization list.

To capture acute beverage-related subjective effects, oral freshness, whole-body freshness, and hunger were assessed immediately before and immediately after the first standardized 200-mL intake (pre-drink vs post-drink). Repeated subjective states (fatigue and enjoyment) and perceived unity (IOS) were assessed across the spectating session at pre-game, halftime (during-game), immediately post-game, and 15 min post-game.

HR was recorded continuously throughout the session. Saliva was collected by passive drool at pre-game, halftime (during-game), immediately post-game, and 15 min post-game for endocrine assays.

### 2.6 Measures

#### Heart-rate recording

HR was recorded using an optical HR sensor (Polar Verity Sense; Polar Electro Oy, Finland) worn on the forearm. The device provided HR values in beats per minute (bpm) at 1-Hz resolution (one sample per second). We analyzed the raw bpm time series without additional preprocessing (e.g., artifact removal or interpolation). For subsequent synchrony analyses, participants’ HR time series were time-aligned to a common game timeline.

#### Interpersonal HR synchrony

Interpersonal HR synchrony was assessed using cross-correlation function (CCF) analysis, following the approach used in our prior spectator-study framework (Matsui et al., 2025). Cross-correlations were computed in R for four predefined segments: first half, halftime, second half, and the 15-min post-game period. Segment boundaries were defined using game event timestamps: first half (tip-off to halftime start), halftime (halftime start to second-half start), second half (second-half start to final buzzer), and 15-min post-game (final buzzer to 15 min thereafter). HR time series were cropped to these intervals prior to analysis.

A temporal time-lag window of ±30 s was applied to identify the maximum correlation between HR time series, matched to the characteristic duration of a basketball play. From each pairwise CCF, we derived two primary indices used in figures: HR synchrony (CCF), defined as the peak cross-correlation coefficient (r_peak), and HR time lag (sec), defined as the absolute peak lag (|lag_peak|), where smaller values indicate tighter near-zero-lag alignment. Dyadic indices were averaged across all pairings to obtain participant-level synchrony scores within each segment.

#### Psychological measures

Subjective states were assessed using visual analog scales (VAS; 0–100 mm) with endpoints labeled 0 = not at all and 100 = extremely. VAS items included oral freshness, whole-body freshness, fatigue, enjoyment, and hunger. Oral freshness, whole-body freshness, and hunger were administered twice around the first standardized intake (pre-drink and immediately post-drink). Fatigue and enjoyment were administered at the four main assessment time points (pre-game, halftime, immediately post-game, 15 min post-game).

Perceived social connection was assessed using the Inclusion of Other in the Self (IOS) scale. Participants rated perceived unity toward (i) home-team players and (ii) home spectators using the standard single-item IOS pictorial measure.

#### Salivary biomarkers (oxytocin, cortisol)

Saliva samples were collected by passive drool at the four main assessment time points (pre-game, halftime, immediately post-game, 15 min post-game). Immediately after collection, samples were placed in a −20°C container and transferred to a −80°C freezer within 3 h for long-term storage until assay. Salivary oxytocin concentrations were quantified using a commercially available enzyme immunoassay kit (Enzo Life Sciences; distributed by Cosmo Bio, Japan) following the manufacturer’s instructions.

Primary oxytocin analyses focused on the three post-baseline time points (halftime, immediately post-game, and 15 min post-game), and pre-game oxytocin is reported separately (Supplementary Fig. S3).

#### Oxytocin–HR synchrony association

We tested association between oxytocin dynamics and post-game preservation of HRsynchrony. Oxytocin change was expressed as a percent-of-baseline index at 15 min post-game relative to halftime: Oxytocin change index (%) = 100 × Oxytocin _{15min post}/ Oxytocin _{halftime} (lower values indicate larger decreases). ΔHR time lag was computed as the change in HR time lag from the first half to 15 min post-game (ΔHR time lag = |lag_peak|_{15min post} − |lag_peak|_{first half}), where smaller values indicate better preservation of near-zero-lag synchrony.

### 2.7 Statistical analysis

All statistical analyses were conducted in R. Outcomes were analyzed using linear mixed-effects models (LMMs) with Group (SW vs PW) as a between-subject factor and Time as a within-subject factor, including the Group × Time interaction. Participant was included as a random intercept to account for repeated measurements.

Time levels differed by outcome based on the measurement schedule. For fatigue, enjoyment, and IOS, Time had four levels (pre-game, halftime, immediately post-game, 15 min post-game). For oral freshness, whole-body freshness, and hunger, Time had two levels (pre-drink and immediately post-drink). For HR synchrony indices, Time had four levels (first half, halftime, second half, 15-min post-game). For salivary oxytocin, Time had four levels (pre-game, halftime, immediately post-game, 15 min post-game); primary oxytocin analyses were conducted on the three post-baseline time points, and pre-game oxytocin was analyzed separately (Supplementary fig. 3).

When the Group × Time interaction was significant, we conducted Bonferroni-corrected pairwise comparisons to characterize between-group differences at specific time points and/or within-group changes across time. Effect sizes were reported as Cohen’s d for pairwise comparisons and partial η² for LMM effects. Statistical significance was set at a two-sided alpha level of 0.05.

To quantify post-game preservation of near-zero-lag synchrony, we computed ΔHR time lag as the change in absolute peak time lag from the first half to 15 min post-game (ΔHR time lag = |lag_peak|_{15min post} − |lag_peak|_{first half}) (smaller values indicate better preservation). We also computed an oxytocin change index at 15 min post-game relative to halftime: Oxytocin change index (%) = 100 × Oxytocin _{15min post}/ Oxytocin _{halftime} (lower values indicate larger decreases). Associations between (i) ΔHR time lag and IOS ratings at 15 min post-game, and (ii) Oxytocin change index (%) and ΔHR time lag, were evaluated using Pearson correlation analyses.

## 3. Results

### 3.1 Acute subjective responses to the first standardized intake

We first examined acute subjective responses to the first standardized beverage intake (200 mL), assessed immediately before and immediately after drinking. Oral freshness, whole-body freshness, fatigue, and hunger were analyzed using mixed models with Group (SW vs. PW), Time (pre-drink vs. post-drink), and the Group × Time interaction (Fig. 2).

**Figure 2.**
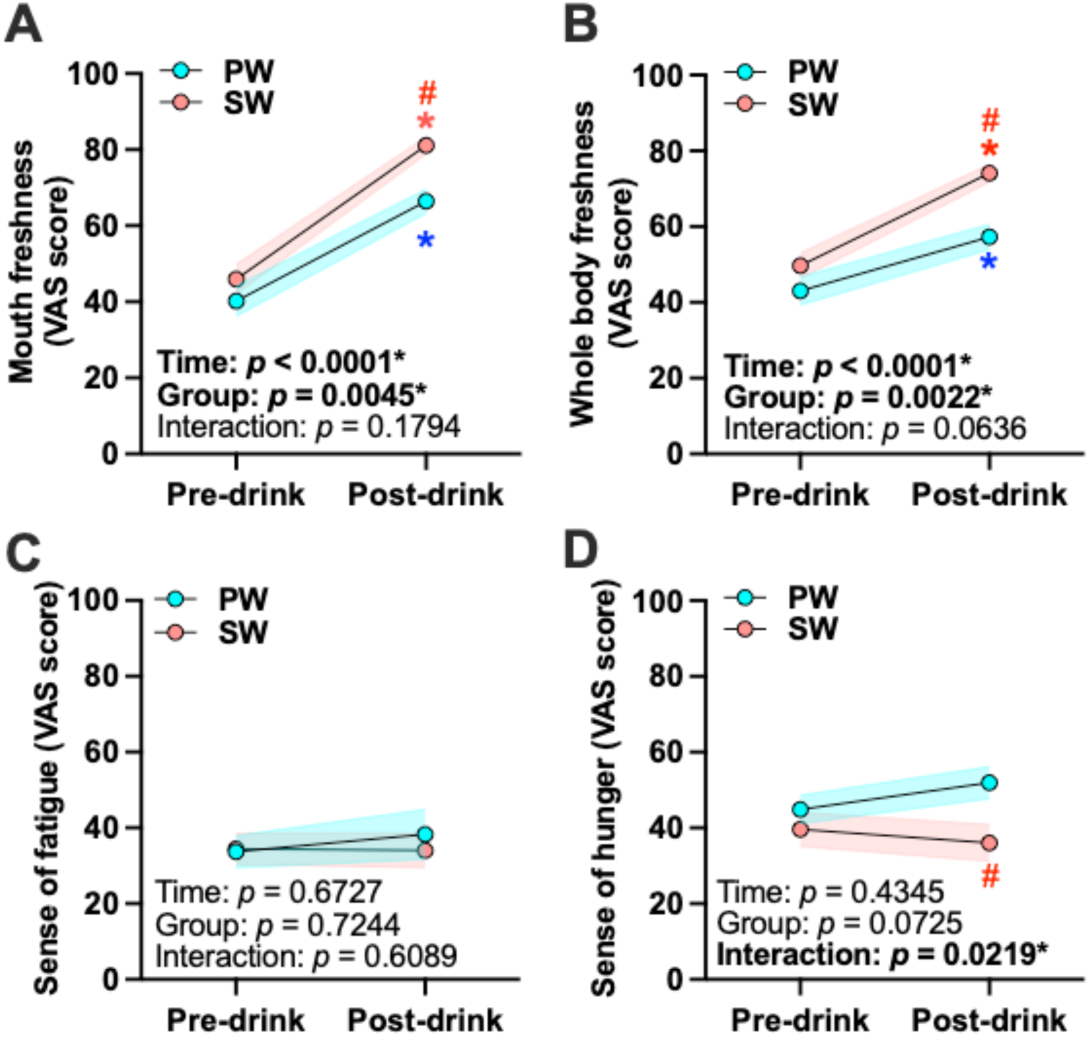
Acute subjective effects of the first standardized intake. VAS ratings assessed immediately before (pre-drink) and after (post-drink) the first standardized 200-mL intake. (A) Oral freshness, (B) whole-body freshness, (C) fatigue, and (D) hunger are shown for PW and SW. Lines indicate group means with shaded bands indicating variability (SEM). Panel annotations report p-values for Time, Group, and the Group × Time interaction from linear mixed-effects models. Where indicated, post-hoc comparisons were performed with Holm–Bonferroni correction. \**p* < 0.05 vs. during game within the same group. #*p* < 0.05 vs. PW group at the indicated time point.

#### Oral freshness

There was a strong main effect of Time (F(1, 78) = 87.45, p < 0.0001, partial η² = 0.529) and a main effect of Group (F(1, 78) = 8.575, p = 0.0045, partial η² = 0.099), whereas the Group × Time interaction was not significant (F(1, 78) = 1.835, p = 0.1794, partial η² = 0.023) (Fig. 2A).

#### Whole-body freshness

Time was significant (F(1, 78) = 51.49, p < 0.0001, partial η² = 0.398) and Group was significant (F(1, 78) = 10.03, p = 0.0022, partial η² = 0.114), whereas the Group × Time interaction did not reach significance (F(1, 78) = 3.541, p = 0.0636, partial η² = 0.043) (Fig. 2B).

#### Fatigue

No effects were detected in the acute fatigue ratings (Time: p = 0.6727; Group: p = 0.7244; Group × Time: p = 0.6089) (Fig. 2C).

#### Hunger

The Group × Time interaction was significant (F(1, 78) = 5.473, p = 0.0219, partial η²= 0.0656), indicating that the pre-to-post change differed by group. The main effects of Time (F(1, 78) = 0.617, p = 0.435, partial η² = 0.00785) and Group (F(1, 78) = 3.315, p = 0.0725, partial η² = 0.0408) were not significant (Fig. 2D).

### 3.2 Subjective state and perceived unity across the session

Fatigue and enjoyment were assessed at four time points (pre-game, halftime, immediately post-game, and 15 min post-game) and analyzed using mixed models with Group, Time, and the Group × Time interaction. IOS-based unity toward home-team players and home spectators was analyzed in the same framework (Fig. 3).

**Figure 3.**
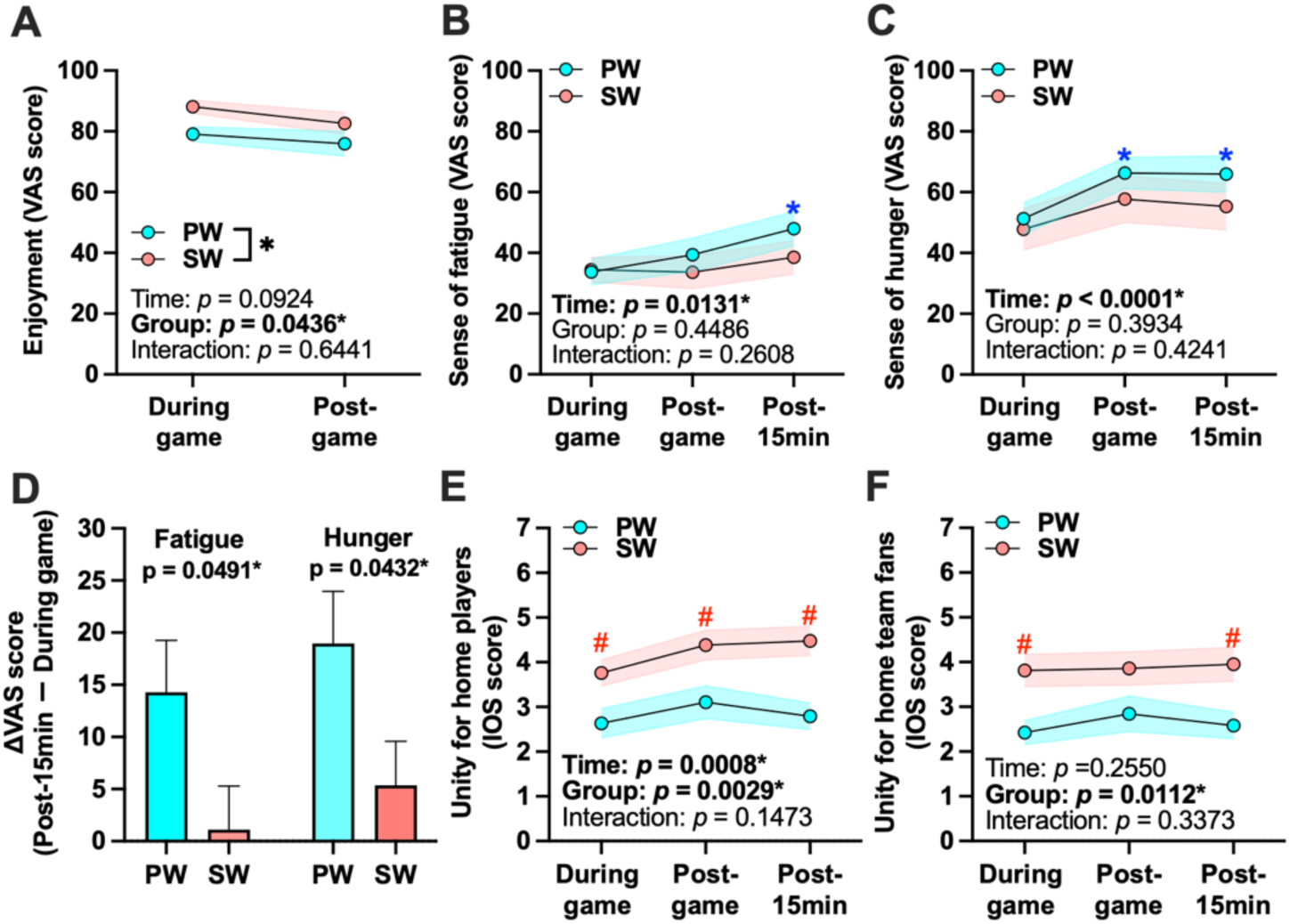
Subjective state and perceived unity during and after the game. Repeated measures of (A) enjoyment, (B) fatigue, and (C) hunger during the game, immediately post-game, and 15 min post-game (as shown) for PW and SW. (D) Change scores (15 min post-game minus during game) for fatigue and hunger are compared between groups. IOS-based perceived unity toward (E) home-team players and (F) home-team fans (home spectators) is shown across time points. Panel annotations report p-values for Time, Group, and Group × Time effects (mixed models) and for between-group comparisons of change scores. Lines indicate group means with shaded bands indicating variability (SEM). Where indicated, post-hoc comparisons were performed with Holm–Bonferroni correction. \**p* < 0.05 vs. during game within the same group. #*p* < 0.05 vs. PW group at the indicated time point.

#### Enjoyment

Enjoyment did not show a significant change over time (F(2, 76) = 5.644, p = 0.0924, partial η² = 0.129), while the Group × Time interaction (F(2, 76) = 1.507, p = 0.6441; partial η² = 0.038). In contrast, a main effect of Group was observed, with the SW group reporting higher enjoyment than the PW group across the session (F(1, 38) = 1.734, p =0.0436, partial η² = 0.1958; Fig. 3A).

#### Sense of fatigue

There was a main effect of Time (F(2, 76) = 4.593, p = 0.0131, partial η² = 0.108), and the Group × Time interaction was not significant (F(2, 76) = 1.368, p = 0.2608, partial η² = 0.0347) and the main effect of Group (F(1, 38) = 0.586, p = 0.4486, partial η² = 0.0152) were not significant (Fig. 3B). In addition, a planned comparison on the change in fatigue from halftime to 15 min post-game differed between groups (Δfatigue_{15min post − halftime}: t(37) = 2.035, p = 0.0491, Cohen’s d = 0.652; Fig. 3D), consistent with attenuated post-game fatigue build-up in the SW group.

#### Hunger

There was a main effect of Time (p < 0.0001), whereas the Group × Time interaction (p = 0.4241) and the main effect of Group (p = 0.3934) were not significant (Fig. 3C). In addition, a planned comparison on the change in hunger from during game to 15 min post-game showed a significant between-group difference, with the SW group exhibiting a lower change score (p = 0.0432; Fig. 3D).

#### Unity toward home-team players

The Group × Time interaction was not significant (F(3, 114) = 4.262, p = 0.1473, partial η² = 0.1008). There were significant main effects of Time (F(3, 114) = 30.79, p = 0.0008, partial η² = 0.4476) and Group (F(1, 38) = 7.966, p = 0.0029, partial η² = 0.1733) (Fig. 3E).

#### Unity toward home spectators

The Group × Time interaction (F(3, 114) = 0.4875, p = 0.3373, partial η² = 0.0127) and a main effect of Time (F(3, 114) = 5.263, p = 0.2550, partial η² = 0.1217) were not significant. Whereas, there was a a main effect of group (F(1, 38) = 7.630, p = 0.0112, partial η² = 0.1672) were significant (Fig. 3F).

### 3.3 Heart-rate responses and interpersonal HR synchrony

As a context check, mean and peak HR increased from the pre-game baseline to the during-game period, accompanied by stronger interpersonal HR synchrony as reflected in higher CCF-based synchrony and shorter HR timelag (Supplementary fig. 1A–D), consistent with elevated arousal and shared engagement during live spectating (Matsui et al., 2025).

We then quantified interpersonal HR synchrony across four predefined segments (first half, halftime, second half, and the 15-min post-game period) using cross-correlation indices within a ±30-s lag window (Fig. 4) Mixed models were used with Group, Time, and the Group × Time interaction.

**Figure 4.**
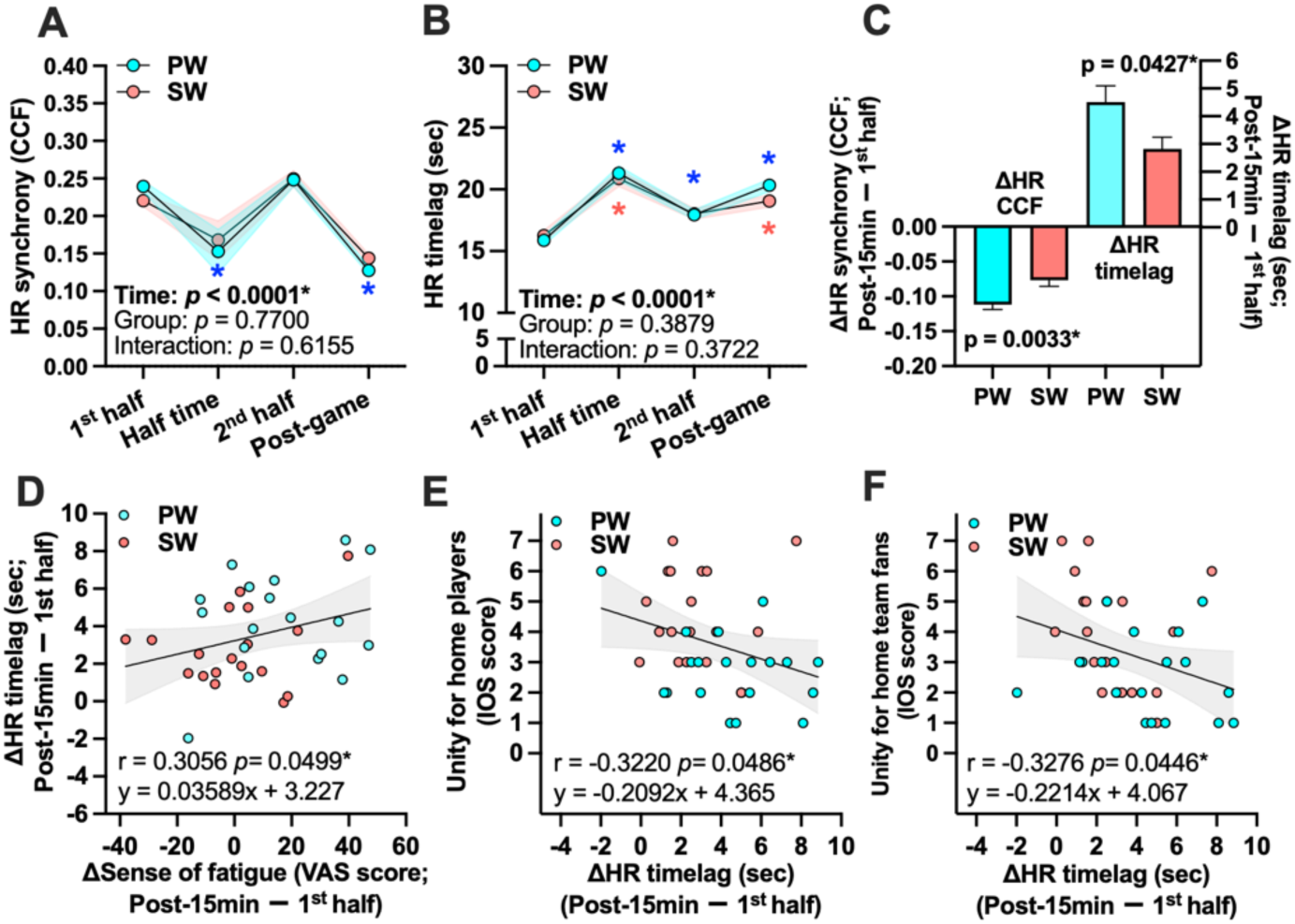
Interpersonal HR synchrony, post-game preservation, and associations with fatigue and unity. Interpersonal HR synchrony, post-game preservation, and associations with fatigue and unity. (A) HR synchrony (CCF) and (B) HR timelag (sec) across segments (first half, halftime, second half, and post-game) for PW and SW. (C) Group comparisons of change scores from the first half to 15 min post-game for ΔHR synchrony (CCF) and ΔHR timelag. Lines indicate group means with shaded bands indicating variability (SEM). Where indicated, post-hoc comparisons were performed with Holm–Bonferroni correction. \**p* < 0.05 vs. during game within the same group. (D) Association between change in fatigue (15 min post-game minus first half) and ΔHR timelag. (E–F) Associations between ΔHR timelag and IOS-based unity at 15 min post-game toward (E) home-team players and (F) home-team fans. Scatter plots show individual data points with regression lines and confidence bands; Pearson’s r and p are reported in panels.

#### Heart rate

Mean and peak HR increased across the spectating session, from the pre-game baseline to in-game and post-game segments (Time: p < 0.0001 for both HRmean and HRpeak). In contrast, beverage group effects were not detected for either HRmean (Group: p = 0.4848; Group × Time: p = 0.4204) or HRpeak (Group: p = 0.5411; Group × Time: p = 0.7452), indicating that beverage condition did not measurably influence mean or peak heart rate (Supplementary fig. 2A, B).

#### HR synchrony (CCF)

For HR-dynamics similarity (HR synchrony (CCF); r_peak), the Group × Time interaction was not significant (F(2.999, 105.0) = 0.9478, p = 0.6155, partial η² = 0.0264), whereas Time showed a strong effect (F(2.999, 105.0) = 245.4, p < 0.0001, partial η² = 0.875) and Group was not significant (F(1, 35) = 0.4985, p = 0.7700, partial η² = 0.0140) (Fig. 4A).

#### HR time lag (sec)

For HR timelag (sec) (|lag_peak|), the Group × Time interaction was not significant (F(3.007, 105.2) = 0.4120, p = 0.3722, partial η² = 0.0116), whereas Time showed a strong effect (F(3.007, 105.2) = 248.6, p < 0.0001, partial η² = 0.877) and Group was not significant (F(1, 35) = 0.3810, p = 0.3879, partial η² = 0.0108) (Fig. 4B).

#### Synchrony preservation

Planned comparisons focusing on synchrony preservation showed that the first-half to post-game change differed between groups for CCF (ΔCCF_{post − first half}: t(36) = 3.149, p = 0.0033, Cohen’s d = 1.02), and for timelag (ΔHR timelag_{post − first half}: t(36) = 2.101, p = 0.0427, Cohen’s d = 0.682) (Fig. 4C).

#### Associations with fatigue change

Change in fatigue (15 min post-game minus during game) was associated with post-game synchrony preservation: Δfatigue was associated with ΔHR synchrony (CCF) (*r* = 0.4777, *p* = 0.0028) and with ΔHR timelag (*r* = 0.3056, *p* = 0.0499; Fig. 4D).

#### Associations with unity

ΔHR timelag (15 min post-game − first half) was negatively associated with IOS-based unity at 15 min post-game toward home-team players (r = −0.3220, p = 0.0486; Fig. 4E) and toward home-team fans (r = −0.3276, p = 0.0446; Fig. 4F), indicating that better near-zero-lag preservation (smaller ΔHR timelag) corresponded to higher perceived unity.

### 3.4 Salivary oxytocin dynamics and associations

#### Oxytocin levels

Salivary oxytocin was analyzed across three post-baseline time points (halftime, immediately post-game, and 15 min post-game) (Fig. 5A). For oxytocin, neither the Group × Time interaction (F(1.722, 51.66) = 2.066, p = 0.1027, partial η² = 0.0644) nor the main effects of Time (F(1.722, 51.66) = 2.461, p = 0.1432, partial η² = 0.0758) and Group (F(1, 30) = 0.804, p = 0.3770, partial η² = 0.0261) were significant. Pre-game oxytocin levels are reported separately in Supplementary figure 3, and baseline oxytocin concentrations did not differ between the PW and SW groups (p = 0.7801).

**Figure 5.**
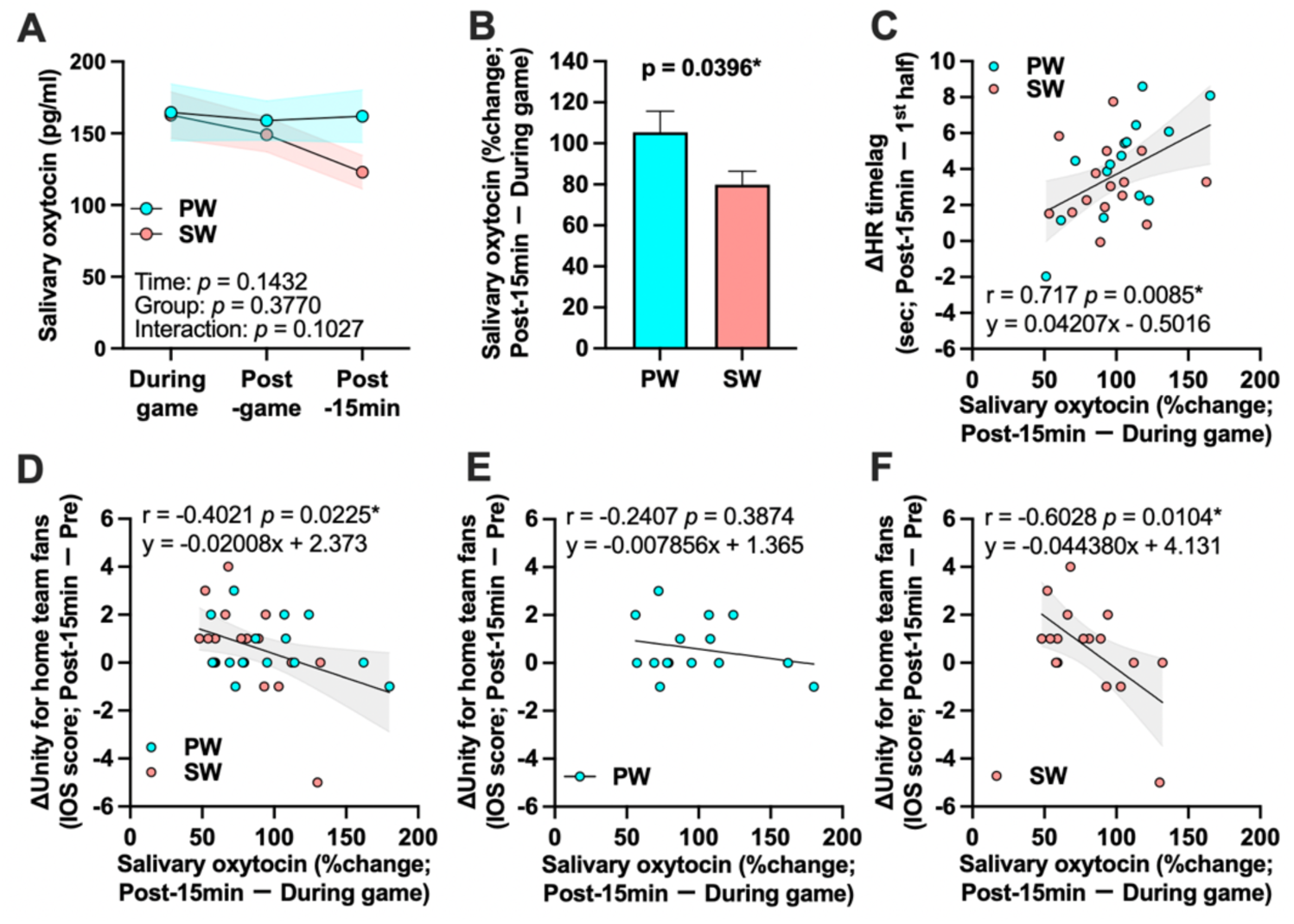
Salivary oxytocin dynamics and their associations with HR synchrony and unity. (A) Salivary oxytocin concentrations during the game (halftime), immediately post-game, and 15 min post-game in PW and SW (model p-values shown). (B) Group comparison of oxytocin change from during game to 15 min post-game (percent change). (C) Association between oxytocin change index and ΔHR timelag (15 min post-game minus first half). (D–F) Associations between oxytocin change and change in IOS-based unity toward home-team fans (15 min post-game minus pre-game), shown for all participants (D), PW only (E), and SW only (F). Regression lines and confidence bands are shown; Pearson’s r and p are reported in panels.

#### Oxytocin change

Oxytocin change was expressed as a percent-of-baseline index at 15 min post-game relative to halftime (Oxytocin change index (%) = 100 × Oxytocin (Post-15min/ halftime); lower values indicate larger decreases) (Fig. 5B). The oxytocin change index was significantly lower in the SW group than in the PW group (p = 0.0396), indicating a larger post-game decrease in salivary oxytocin in the SW group.

#### Association with synchrony preservation

Oxytocin change index (%) was positively associated with ΔHR timelag (Post-15min − first half) (r = 0.717, p = 0.0085; Fig. 5C), such that larger oxytocin decreases (lower percent-of-baseline values) were associated with Δ HR timelag, suggesting greater preservation of HR synchrony.

#### Associations with unity change

Oxytocin change index (%) was associated with change in IOS-based unity toward home-team fans (15 min post-game minus pre-game) when pooling participants (r = −0.4021, p = 0.0225; Fig. 5D). Stratified analyses showed no association in PW (r = −0.2407, p = 0.3874; Fig. 5E) and a significant association in SW (r = −0.6028, p = 0.0104; Fig. 5F).

## 4. Discussion

In an individually randomized field experiment during a collegiate women’s basketball home game, we tested the hypothesis that co-consuming sparkling water attenuates hunger and fatigue and enhances enjoyment and perceived unity, potentially via post-game HR synchrony linked to salivary oxytocin dynamics. Consistent with this model, sparkling water improved acute subjective state (freshness and hunger; Fig. 2), and enjoyment was higher overall in the sparkling-water group (Fig. 3A). Planned comparisons indicated a smaller post-game rise in fatigue and a lower change score in hunger (Fig. 3B-D). Sparkling water was also associated with better preservation of post-game synchrony based on planned contrasts for HR synchrony (CCF) and HR timelag (sec). In parallel, HR synchrony preservation was associated with both fatigue change and perceived unity toward players and fans (Fig. 4). Finally, the oxytocin change index differed between groups and was associated with ΔHR timelag and unity toward fellow fans (Fig. 5), providing convergent endocrine support for the coupling pathway. Together, these findings support an alcohol-free shared-drink ritual as a low-burden approach to strengthening socio-physiological coupling and perceived unity in a naturalistic crowd.

### 4.1 Sparkling water shifts subjective state supporting sustained engagement

Sparkling-water co-consumption produced immediate changes in subjective state that are plausibly relevant to spectatorship engagement. Around the first standardized intake, sparkling water increased oral and whole-body freshness and yielded a larger reduction in hunger relative to plain water (Fig. 2A–B, Fig. 2D). These acute effects are consistent with the notion that oral CO₂ stimulation engages trigeminal and oral somatosensory pathways that shape perceived freshness and appetite-related sensations (Suzuki et al., 2017). Mechanistically, carbonated water can excite oral nociceptive pathways via a carbonic anhydrase dependent process, providing a neurobiological route by which carbonation can elicit salient oral sensations that shift subjective state (Dessirier et al., 2000; Simons et al., 1999). Acute fatigue did not differ between groups (Fig. 2C), suggesting that immediate fatigue sensations were not strongly altered by the first intake.

Across the spectating session, fatigue increased over time, and a planned comparison indicated that the post-game increase in fatigue was smaller in the sparkling-water group (Fig. 3B, D). Also, the change in hunger from during game to 15 min post-game was lower in the sparkling-water group (Fig. 3C–D). These patterns are consistent with an engagement-maintenance account in which refreshed subjective state and attenuated fatigue and hunger accumulation help sustain a shared spectator state beyond the final buzzer. This interpretation aligns with our prior esports findings showing that sparkling water reduced fatigue and hunger and increased enjoyment during prolonged play (Takahashi et al., 2025). Consistent with this prior work, enjoyment did not change significantly over time but was higher overall in the SW group (Fig. 3A), suggesting a modest upward shift in affective tone alongside attenuated fatigue accumulation

More broadly, oral sensory stimulation can influence central processing linked to motivation and performance, and oral sensing paradigms have been shown to modulate brain activity in reward-related regions including orbitofrontal cortex (Kosugi et al., 2024; Chambers et al., 2009; Kringelbach, 2005). This affective shift is compatible with a reward/valuation account in which oral sensory stimulation and reduced fatigue help sustain positive engagement during collective events, thereby elevating overall enjoyment. Together, these state effects provide a plausible experiential route through which a shared-drink ritual could contribute to preserved post-game socio-physiological coupling.

### 4.2 Post-game synchrony preservation predicts perceived unity

Interpersonal physiological synchrony has been proposed as an objective signature of moment-to-moment social coupling during shared experiences, reflecting coordinated attentional and affective dynamics across individuals (Palumbo et al., 2017). In the present field setting, sparkling water was associated with reduced post-game decay in coupling. Although the Group × Time interaction was not consistently supported across synchrony indices, planned contrasts indicated that the first-half to post-game change differed between groups for both HR synchrony (CCF) and HR timelag (sec) (Fig. 4C–D). These patterns suggest that a simple add-on ritual can buffer coupling decay during the transition from the structured game context to the immediate post-game period.

Importantly, coupling preservation was directly related to bonding-relevant experience. ΔHR timelag was negatively associated with IOS-based unity at 15 min post-game toward home-team players (Fig. 4E) and toward home-team fans (Fig. 4F), indicating that better HR synchrony preservation (smaller ΔHR timelag) corresponded to higher perceived unity. Related work in other collective contexts has documented physiological synchrony among participants and linked synchrony to affiliative outcomes and cooperative success (Behrens et al., 2020; Konvalinka et al., 2011). Together, our results extend this literature by showing that synchrony preservation in a naturalistic crowd is sensitive to a feasible co-consumption manipulation and is meaningfully related to perceived unity.

### 4.3 Why sparkling-water co-consumption preserves post-game social connection

#### State pathway: oral CO₂ stimulation and engagement

Sparkling-water co-consumption may preserve post-game HR synchrony by stabilizing an engagement-relevant subjective state during and after the game. Carbonation is a salient oral stimulus, and mechanistic work indicates that CO₂ can engage trigeminally mediated oral sensation through carbonic anhydrase–dependent processes (Suzuki et al., 2017; Dessirier et al., 2000; Simons et al., 1999). These mechanisms provide a plausible route to the acute increases in perceived freshness and shifts in appetite-related sensations observed around intake (Fig. 2A–B, Fig. 2D).

In the present setting, these state shifts, together with attenuated post-game fatigue and hunger build-up (Fig. 3B–D), are consistent with an account in which refreshed spectators sustain shared attention and affective engagement beyond the final buzzer, thereby limiting drift in autonomic dynamics after the focal event including esports (Takahashi et al., 2025). Oral sensory stimulation can also influence central processing linked to motivation and valuation, including orbitofrontal regions (Chambers et al., 2009; Kringelbach, 2005). Reports that carbonated water modulates prefrontal hemodynamics near orbitofrontal cortex provide an additional rationale for why sparkling water attenuated post-game fatigue and elevated overall enjoyment(Kosugi et al., 2024), supporting engagement during extended experiences (Fig. 3A, Fig. 3D).

#### Coordination pathway: micro-ritual co-consumption

A second, complementary mechanism concerns coordinated micro-ritual behavior. The manipulation involved synchronized co-consumption timing, and brief repeated coordination can strengthen affiliation and prosociality (Mogan et al., 2017; Reddish et al., 2013). In collective settings, synchrony and coordinated arousal have been linked to increased cohesion and cooperative tendencies (Jackson et al., 2018; Konvalinka et al., 2011). The present pattern, buffering post-game decay rather than elevating synchrony uniformly, is compatible with the idea that synchronized drinking helps carry a shared state into the immediate post-game period, when coupling might otherwise drift.

Coordinated co-consumption could plausibly promote affiliation in both groups because synchronized drinking constitutes a shared micro-ritual regardless of beverage type, and similar consumption has been shown to increase trust and cooperation (Woolley & Fishbach, 2017). Behavioral synchrony more broadly increases cooperation and perceived bonding across contexts (Mogan et al., 2017; Reddish et al., 2013). In the present study, synchrony preservation was stronger in the sparkling-water group and was associated with fatigue change and perceived unity (Fig. 4C–F), consistent with a synergistic account in which sparkling water strengthens the state pathway (reduced hunger/fatigue; Fig. 3D), and thereby allows coordinated co-consumption to more effectively carry coupling into the post-game period (Jackson et al., 2018; Tarr et al., 2015). This interpretation is consistent with the observed association between smaller post-game fatigue increases and better synchrony preservation (Fig. 4D).

#### Oxytocin dynamics at the intersection of state and coordination

Endocrine dynamics provided convergent support for the coupling pathway. The oxytocin change index (%) was lower in the sparkling-water group (Fig. 5B; lower values indicate larger decreases), and oxytocin change index (%) was positively associated with ΔHR timelag (Fig. 5C), such that larger oxytocin decreases were associated with smaller increases in HR timelag and thus greater near-zero-lag preservation. Oxytocin change index (%) was also associated with changes in unity toward home-team fans in the pooled sample and in the sparkling-water group (Fig. 5D–F), providing an endocrine link to bonding-relevant outcomes.

Although peripheral and salivary oxytocin measures have known interpretational and analytical limitations, this pattern is compatible with the possibility that oxytocin-related processes were engaged during spectatorship, with lower salivary concentrations at sampling reflecting transient release followed by rapid clearance and/or receptor-level regulation rather than reduced involvement (Tabak et al., 2023; Quintana et al., 2018; Lefevre et al., 2017; Modi et al., 2016; Smith et al., 2006). Consistent with the idea that decreases in peripheral oxytocin are not necessarily inconsistent with bonding, salivary oxytocin has been reported to decrease in socially engaging group activities while self-reported social connectedness improves (Bowling et al., 2022). Moreover, inverse associations between peripheral oxytocin levels or reactivity and empathy-related performance have been reported in some contexts, underscoring that directionality can vary by population, task, and timing (Papasteri et al., 2020; Montag et al., 2020). Finally, oxytocin change index was associated with changes in unity toward home-team fans in the pooled sample and in the sparkling-water group (Fig. 5D–F), providing additional linkage between endocrine dynamics and bonding-relevant outcomes during spectatorship.

A complementary interpretation is that oxytocin-linked processes may also influence the affective dimension of collective experience, thereby supporting synergy between the state pathway and coordination pathway. Preclinical evidence indicates that oxytocin can modulate mesolimbic reward circuitry via the ventral tegmental area (VTA) dopamine system, and bidirectional oxytocin–dopamine interactions have been proposed as a substrate through which endocrine dynamics could align with reward/valuation processes and prosocial engagement (Hung et al., 2017; Baskerville & Douglas, 2010). In the present study, this perspective is consistent with a reciprocal account in which more positive engagement supports sustained shared attention and coupling after the game (Fig. 3A), while preserved coupling and unity further enhance the subjective experience (Fig. 4E–F), with oxytocin dynamics varying alongside both sides of this loop (Fig. 5B–F).

### 4.4 Limitations and future directions

Several limitations qualify the interpretation and generalizability of our findings. First, the study was conducted in a single basketball game at one venue, so replications across games, sports, and crowd compositions are needed to assess robustness and boundary conditions. Second, participant blinding was not feasible because sparkling water has distinctive sensory properties; expectancy effects may influence subjective reports and behavior, motivating explicit expectancy measures and stronger control beverages where feasible. Third, HR was measured using a forearm-worn PPG device at 1 Hz and analyzed as raw bpm time series; future work could pair PPG with ECG-based recordings and apply artifact-aware preprocessing to strengthen inference. Fourth, HR synchrony in naturalistic settings can reflect shared stimulus structure in addition to interaction-driven coupling; incorporating behavioral observation and analytical approaches that dissociate shared-input effects from interpersonal coordination would sharpen mechanistic conclusions. Fifth, ad libitum intake after standardized doses was not quantified and could have contributed to comfort or subjective state differences, motivating protocols that record intake volume. Finally, matchday drinking is often alcohol-centered, and alcohol availability at sporting events is discussed in relation to alcohol-related problems and venue safety or public health concerns (Ruehlmann et al., 2023; Lenk et al., 2010; Nelson & Wechsler, 2003). Future field trials should test substitution more directly by measuring alcohol use during spectating and comparing sparkling water with alcohol and other non-alcoholic controls.

## 5. Conclusion

This study tested whether a simple, scalable co-consumption ritual, drinking sparkling water together, strengthens bonding-relevant outcomes during live sport spectatorship. Sparkling water improved subjective state (freshness, hunger, and enjoyment) and attenuated post-game fatigue, and it was associated with better preservation of post-game HR synchrony and higher IOS-based perceived unity. Oxytocin dynamics were associated with synchrony preservation, consistent with a model linking engagement-related state, socio-physiological coupling, and bonding in a naturalistic crowd. Together, these findings highlight an alcohol-free shared-drink ritual as a low-burden candidate for enhancing social connection during and after live sport events.

## Data availability

The data supporting the findings of this study are available from the corresponding author upon request.

## Author contribution

T.M., W.K., and S.M. conceived and designed the study. T.M., T.Y., S.D., S.T., Y.T., and R.T. recruited participants, collected the data, conducted data analysis and interpreted data. T.M. and T.Y. drafted the manuscript. T.M., T.Y., W.K., and S.M. edited and revised the manuscript. All authors approved the final version.

---

**Supplementary figure 1.**
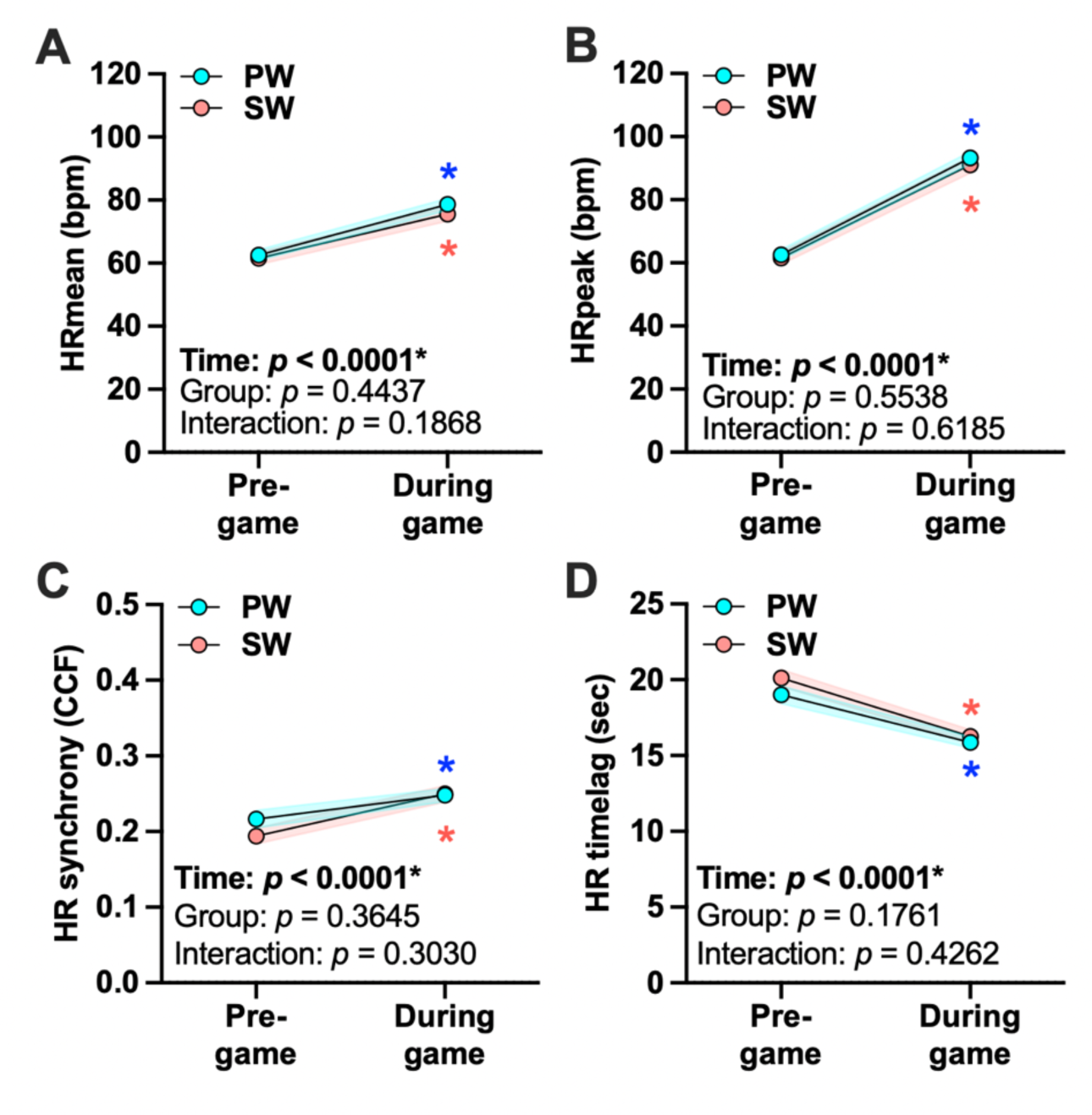
Pre-game versus during-game changes in heart rate and its synchrony. (A) Mean heart rate, (B) peak heart rate, (C) CCF-based HR synchrony, and (D) HR time lag are compared between pre-game and during-game periods for PW and SW. Panel annotations report p-values for Time, Group, and Group × Time effects. Lines indicate group means with shaded bands indicating variability (SEM). Where indicated, post-hoc comparisons were performed with Holm–Bonferroni correction. \**p* < 0.05 vs. during game within the same group.

**Supplementary figure 2.**
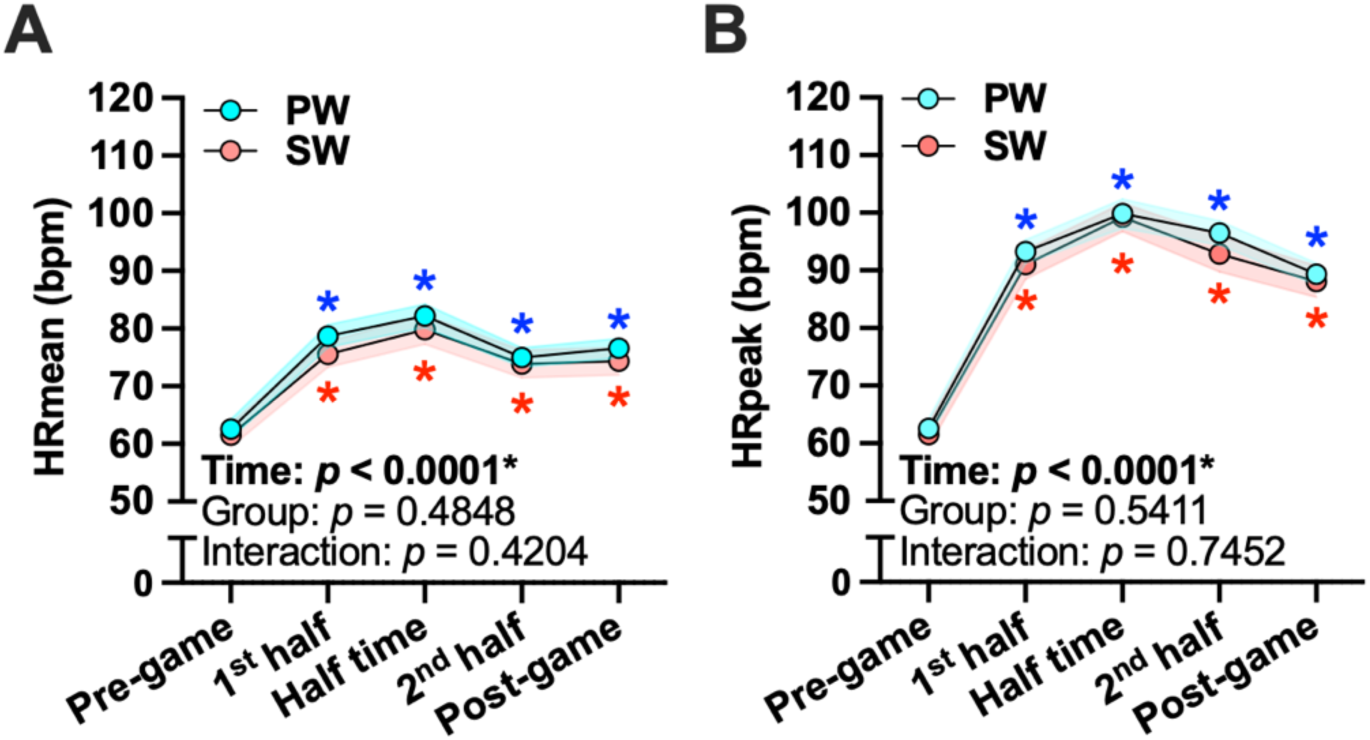
Mean and peak heart rate across game segments. (A) HRmean and (B) HRpeak across segments (pre-game, first half, halftime, second half, and post-game) for PW and SW. Panel annotations report p-values for Time, Group, and Group × Time effects (mixed models). Lines indicate group means with shaded bands indicating variability (SEM).

**Supplementary figure 3.**
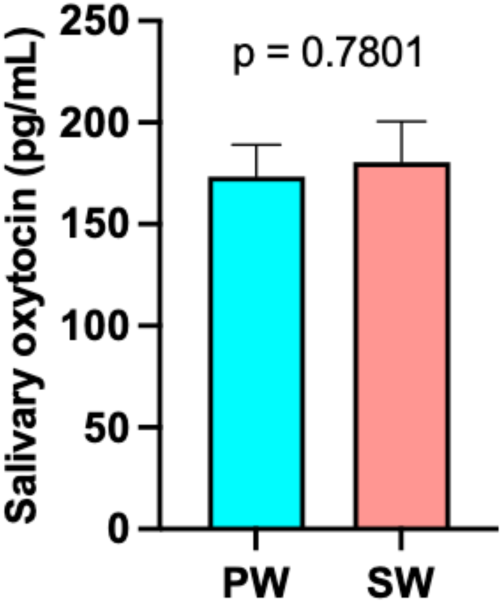
Salivary oxytocin at the pre-game baseline. Salivary oxytocin concentrations at the pre-game baseline are shown for PW and SW. Data are presented as mean ± SEM. The p-value shown in the panel was obtained from an independent-samples t-test (two-sided).

